# Pre-Clinical Evaluation of a Novel Immunomodulator: a potential immunotherapy for coronaviral disease

**DOI:** 10.1101/2025.04.01.646433

**Authors:** Irene Lee, Amar Desai, Akshay Patil, Yan Xu, Kelley Pozza-Adams, Anthony Berdis

## Abstract

At present, no unifying animal study to quantify the effect of PI on innate immunity in animals has been reported. This study quantifies the effect of PI in mice and THP-1 cells using a multi-disciplinary approach employing flow cytometry to measure immune function and mass spectroscopy to identify metabolomic profiles of treated cells. For proof of principle, a case study was conducted to examine the potential benefit of administering PI to improve outcomes of a feline leukemia-positive cat that also contracted the effusive form of FIP. Our results indicate that quantifiable cellular and metabolic markers result from PI treatment and can be used to establish a platform measuring the efficacy of PI in modulating the innate immune system.

## Introduction

Coronaviruses (CoVs) are a family of enveloped viruses containing a positive-sense RNA genome that infect a broad range of vertebrates including mammals and birds. ^1–4^ While over 100 species of CoVs have been identified,^5, 6^ only seven CoV strains have been isolated from humans.^7–10^ Of these, four strains (HCoV-229E, HCoV-NL63, HCoV-HKU1, and HCoV-OC43) usually produce mild upper respiratory infections and account for an estimated 15-30% of all reported cases of the “common cold.” ^11^ However, the three most recently identified human CoV strains pose significantly greater health threats as they produce more serious respiratory infections. These include severe acute respiratory syndrome (SARS-CoV), identified in 2002; Middle East respiratory syndrome (MERS-CoV), identified in 2012; and SARS-CoV-2, identified in 2019.^12–19^ This most recent HCoV is by far the most significant as it has infected more than 700 million people and caused the death of more than 7 million people worldwide. ^20–22^

In addition to humans, CoVs infect a large number of mammalian and avian species to cause diseases in companion animals (cats and dogs) and livestock (cattle, pigs, and chickens). ^2, 4, 19, 23^ In this study, we used feline coronavirus (FCoV) as a model for human CoVs due to the striking similarities between the two. At the cellular level, SARS-CoV-2 and FCoV possess a large (∼30 kb) positive-sense, single-stranded RNA genome containing 11 open reading frames (ORFs). Four ORFs encode structural proteins including the spike (S), envelope (E), membrane (M), and nucleocapsid (N) proteins. Five ORFs encode the nonstructural proteins 3a, 3b, 3c, 7a, and 7b.^24^ The other two major ORFs encode the RNA-dependent RNA polymerase (RdRp) and helicase which function as the viral replicase.

There are also several relevant clinical similarities between SARS-CoV-2 and FCoV infections. For instance, humans infected with SARS-CoV-2 and felines infected with FCoV can be asymptomatic or display mild symptoms including fever, diarrhea, dehydration, and sneezing/coughing. In felines, however, persistent infection leads to mutational events that transform FCoV into a highly virulent strain called feline infectious peritonitis virus (FIPV).^25–27 28–30^ FIPV is macrophage tropic and is believed to cause aberrant cytokine and/or chemokine expression which causes lymphocyte depletion. This culminates in a highly fatal disease called feline infectious peritonitis (FIP) and is similar to the “cytokine storm” reported in COVID-19 patients that causes life-threatening symptoms such as pulmonary infiltrates and cardiovascular shock. ^64–66^

To date, there are few antiviral agents that can combat CoV infections. One notable exception is the nucleoside analog, Remdesivir (Gilead Science), which received emergency FDA approval in late 2020 to treat certain patients infected with SARS-CoV-2.^23,24^ When tested in cell culture systems, Remdesivir showed efficacy toward inhibiting SARS-CoV and MERS-CoV replication. A recent NIH clinical trial showed that Remdesivir treatment in SARS-CoV-2 infected patients reduced the median time to recovery from 11 days to 15 days.^31^ Unfortunately, Remdesivir did not significantly improve mortality in patients infected SARS-CoV-2. Another nucleoside analog, molnupiravir, is effective in treating mild to moderate COVID-19 in non-hospitalized patients. However, its effectiveness in treating late-stage or severe COVID-19 patients is unclear.^32^ Furthermore, molnupiravir does not appear to reduce the risk of mortality and hospitalization of vaccinated patients. These limitations highlight an urgent need to develop innovative therapeutic strategies and agents to treat severe CoV infections. Interestingly, remdesivir and molnupiravir have been successfully used to treat FIP in non-immune compromised felines within a treatment window of about 84 days, which is much longer than the treatment window for COVID-19 in humans.^33^

An alternative to nucleoside analogs to inhibit viral replication is to develop therapeutic strategies to increase immune function. One attractive candidate is PI (VetImmune), which is a USDA-approved and commercially available small-molecule used in veterinary medicine to treat feline rhinotracheitis (FRV).^34^ More recently, PI has been repositioned as a health supplement for boosting innate immunity in both felines and canines. PI is also used “off-label” to treat felines infected with feline leukemia virus (FeLV, I. Lee unpublished results) and non-effusive FIP.^35–37^ FeLV is an oncornavirus (RNA virus) of the family Retroviridae.^38^ The median survival time for cats infected with FeLV is approximately 2.5 years. In this case, morality is typically caused by severe anemia, abnormal white blood cell counts, and compromised bone marrow function. Felines suffering from FIP and FeLV display similar hematological abnormalities and dysfunction. In addition, FIP induces a globally dysregulated inflammatory response that destroys several major organs, including the lungs, kidneys, and the central nervous system.^39^

In contrast to the nucleoside analogs Remdesivir and Molnupiravir, PI is a chemical mixture of phosphorylated polyisoprenols containing three trans- and several cis-isoprene units (Figure 1). The antiviral efficacy of PI has been demonstrated in felines through analyses of clinically relevant data monitoring changes in the appetite/body weight, blood chemical profiles, and complete blood count values of infected felines.^35–37^ This information shows PI can cause remission of the non-effusive form of FIP. Unfortunately, this clinical-based evidence does not elucidate the mechanism of action of PI. One cell-based study using the human cell line THP-1 showed a modest upregulation in TNFα levels within hours after PI treatment.^40^ Flow-cytometry analyses detected an elevated level of Cd11b in whole blood and isolated monocytes in feline inoculated with FRV and treated with PI over 3 days. Based on these data, the hypothesis is that PI combats viral pathogenesis by reprogramming the cell-mediated innate immune response.

**Figure 1.**
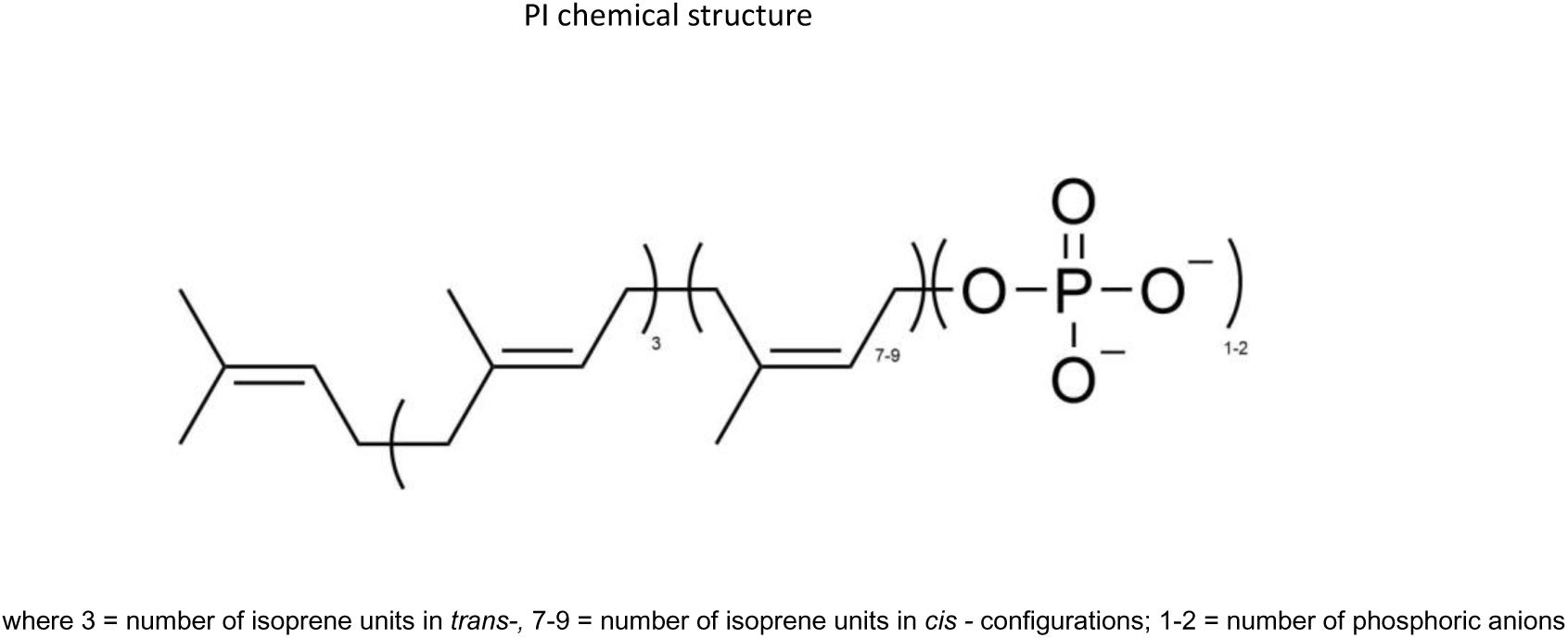

Since the signs of FIP resemble those found in human COVID-19,^41^ we propose that PI can potentially treat COVID and possibly other viral infections in humans. PI could synergize with an antiviral agent such as Molnupiravir in treating viral infections by directing the host’s immune system to actively remove the chemically inactivated viruses. However more rigorous quantitative characterization of the cellular pathways modulated by PI must be performed to reposition PI as a human therapeutic agent. This study bridges this gap in knowledge using a pre-clinical mouse model combined with mammalian cell culture systems to characterize immunological outcomes in response to PI treatment.

## Materials and Methods

### Materials

Phosphate-buffered saline (PBS), antibiotic and antifungal agents, amphotericin, propidium iodide, PrestoBlue, DAPI, Alexa Fluor 488, and apoptosis assay kit containing Alexa Fluor 488-labeled Annexin V were from Invitrogen. Polyprenyl immunostimulant (PI) was supplied from SASS & SASS, Inc. as a solid and dissolved in HPLC grade n-butanol as a 10 mg/mL stock. LC-MS grade acetonitrile and methanol were purchased from Thermo Fischer Scientific (Waltham, MA, USA) for metabolomic studies. Pure acetic acid (99.8) was purchased from Acros Organics, and Deionized water with a resistance of 18.2 MΩ was prepared using a Barnstead GenPure xCAD ultrapure water system from Thermo Fisher Scientific (Waltham, MA, USA). Ammonium acetate (LC-MS grade) was obtained from Sigma-Aldrich (St. Louis, MO, USA). Chloroform was purchased from Sigma-Aldrich. Prostaglandin E2-d4 (PGE2-d4) standard solution (500 μg/mL in methyl acetate, (>99% purity) was purchased from Cayman Chemical (Ann Arbor, MI, USA) and used as the internal standard (IS) master stock. THP-1 cells were obtained from ATCC

### Methods

#### Cell culture

THP-1 cells were cultured in a humidified atmosphere of 5% CO2 at 37 °C. Cells were grown in RPMI (Sigma) supplemented with 10% fetal bovine serum (FBS) (Biowest), 1.0% Penicillin Streptomycin (Gibco) at 37 °C with 5.0% CO2. Since this cell line was obtained from ATCC, informed consent was not required. Cells were routinely authenticated based on morphology and growth characteristics. All cells were expanded and then frozen at low passage (passages 2-5) within 2 weeks after the receipt of the original stocks. All cells used for experiments were between passages 6 and 12. Cell lines were tested for mycoplasma after each thaw or every 4 weeks when grown in culture. Mycoplasma infection was detected using the MycoAlert Mycoplasma Detection kit from Lonza (Walkersville, MD).

#### Cell viability assays

THP-1 cells were plated at an initial density of 200,000 cells/mL. PI was added to wells in a dose-dependent manner (1−100 mg/mL). In all experiments, the final concentration of the co-solvent, n-butanol, was maintained at 0.1%. Cells were treated and assessed for viability for time periods ranging from 2 - 72 hr. Cell viability was assessed using a Muse Cell Count (EMD Millipore) and via Cell-Titer blue assays. ED50 values for PI against THP-1 cells were obtained using a non-linear regression curve fit of the data to Equation 1.

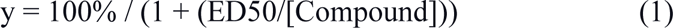

#### Apoptosis Measurements

THP-1 cells were plated at an initial density of 200,000 cells/mL. Cells were treated with variable concentrations of PI, as described above. At variable time intervals (2-72 hours), cells were harvested by centrifugation, washed in PBS, and re-suspended in 100 mL of binding buffer containing 5 mM of Annexin V-Alexa Fluor 488 conjugate. Cells were treated with 1 mg/ mL propidium iodide and incubated at room temperature for 15 min, followed by flow cytometry analysis. Cells were analyzed using a Muse Cell analyzer. 15,000-gated events were observed for each sample.

### Animal Studies

#### CWRU Animals

Animals were housed in the AAALAC-accredited facilities in the CWRU School of Medicine. Experimental procedures were approved by the CWRU Institutional Animal Care and Use Committee (IACUC) in accordance with approved IACUC protocols (2019-0065). Mice were housed in standard microisolator cages and maintained on a defined, irradiated diet and autoclaved water. Experimental procedures used in this arm of animal experiments are based on existing dosage and treatment regimens used to treat felines suffering from FIP (1-4). Mice were treated with 3 mg/kg PI once daily by intraperitoneal (IP) injection for 3 consecutive days. After the final PI treatment (day 3), the mice were euthanized at 8 hours after the final treatment. All animals were observed daily for signs of illness.

#### SASS & SASS Animals

Husbandry and experimental procedures were approved by the Company’s Institutional Animal Care and Use Committee (IACUC). Eight females over one year old CD-1 retired breeders (Charles Rivers lab) that were previously used for toxicity study of PI were administered with 3 mg/Kg of PI.

#### Complete Blood Count Analysis

Peripheral blood was collected into Microtainer EDTA tubes (Becton-Dickinson) by submandibular cheek puncture. Blood counts were analyzed using a Hemavet 950 FS hematology analyzer.

#### Quantification of BM and splenic immune cell populations

Bone marrow cells were obtained by flushing hindlimb bones, and splenocytes were obtained by mincing spleens. Cells were stained with antibodies against CD45R/B220 (RA3-6B2), CD11b (M1/70), CD3e (500A2),, B220(RA3-6B2), CD4 (RM4-5), CD8 (53-6.7), Ly-6G and Ly6C (RB6-8C5), TER-119 (TER-119), Ly-6A/E (D7), CD117 (2B8), F4/80 (Cl:A3-1), CD61 (2C9.G2), and data was acquired on an LSRII flow cytometer (BD Biosciences). Analysis was performed on FlowJo software (TreeStar).

#### Statistical Analysis

All values were tabulated graphically with error bars corresponding to the standard error of the mean. Analysis was performed using GraphPad Prism software. Unless otherwise noted, an unpaired two-tailed Student’s t-test was used to compare treatment groups.

### Metabolomic Studies using Mass Spectroscopy

#### Internal Standard (IS) solution preparation

The working stock solution of PGE2-d4 of 10.0 μg/mL was prepared by diluting 20.0 μL of the 500 μg/mL with 980 μL of diluting solvent methanol/acetonitrile/water (2:2:1).

#### Cell sample preparation

Metabolomic studies with THP-1 were performed using three different treatment regimens corresponding to (untreated cells), treatment with n-butanol (vehicle), and treatment with 75 ug/mL PI for 24 hours, the same condition used in the THP-1 flow cytometry study. After 24 hours, cells were harvested by centrifugation and the resulting cell pellet was suspended in 1.0 mL deionized water and sonicated for 30 seconds to lyse the cells. The protein concentration was measured using the Pierce BCA protein assay kit (Thermo Fischer Scientific) to normalize the cell growth factors across different conditions. The final protein concentration was adjusted to 100 μg/mL by adding an appropriate volume of deionized water. The metabolites of the cell samples were extracted using liquid-liquid extraction by modifying the Folch method. In detail, 1.00 mL of each cell lysate of every experimental condition was added with a 3.00 mL extraction solvent of chloroform/methanol (1/1), and the mixture was vortexed vigorously for 2.00 minutes and placed in an ice bath for 15.0 minutes. After the phase separation, the mixture was centrifuged at 2000 x g at 4C for 15.0 minutes, and the lower organic layer was carefully aspirated without disturbing the protein disc in the center and collected in a fresh culture tube. The remaining aqueous phase and protein disc was added with 3.00 mL chloroform/methanol (2/1), and the mixture was vortexed for 2.00 minutes and incubated in an ice bath for 15.0 minutes. Again, the mixture was centrifuged at 2000 x g at 4C for 15.0 minutes, and the lower organic phase was aspirated and mixed with the previous aspirated aqueous phase and mixed. This organic layer was evaporated under a nitrogen stream using the N-EVAP TM 111 nitrogen evaporator from Organomation (West Berlin, Massachusetts, USA). Once the samples were dried, the internal standard solution and reconstitution solvent (acetonitrile/methanol/water with a ratio of 2:1:1) were added, making the final internal standard concentration at 2.00 μg/mL.

Plasma samples prepared from the whole blood of mice: The plasma samples under three experimental conditions (untreated), vehicle (treated with placebo), and treated (treated with PI) were subjected to protein precipitation for extraction of metabolites. The extraction solvent acetonitrile/methanol (2:1) volume of 960 μL was added to 240 μL of plasma. The mixture was vigorously vortexed for 2.00 minutes and incubated overnight at −20C to enhance the precipitation of proteins. The following day, the samples were centrifuged at 13,000 g for 15.0 minutes at 4C, and the supernatant was collected in fresh borosilicate glass culture tubes. The supernatant was dried under a stream of nitrogen and reconstituted with reconstitution solvent (acetonitrile/methanol/water with a ratio of 2:1:1) and internal standard solution ((IS, see above for preparation method, final concentration 2.00 μg/mL).

#### UHPLC-QTOF/MS analysis

The data was acquired on Agilent’s Infinity II 1290 liquid chromatography unit with a binary pump, a degasser, a multisampler, and a column oven compartment coupled with a 6545-quadrupole time-of-flight mass spectrometer. The data for cell and plasma samples were acquired in both positive and negative electrospray ionization (ESI) modes, using a mobile phase pair containing solvent A as 5 mM ammonium acetate aqueous solution with 0.1 % acetic acid and solvent B as acetonitrile/methanol (80:20) with 5 mM ammonium acetate and 0.1% acetic acid with a gradient elution profile as follows; follows; 0.00-1.00 min (40% B), 12.0 min (75% B), 20.0 min (85% B), 28.0-38.0 min (100% B), 40.0 min (75% B), returning to 40.0-45.0 min (40% B). Each chromatographic run included a 10-minute column pre-equilibration at initial conditions (40% B). The chromatographic separation was carried out using Waters XSelect HSS T3 (2.1 x 150 mm, 2.5 μm) analytical column at 35C with a sample volume of 5 μL. Each biological sample was injected four times as technical replicates in both ESI modes.

#### Data processing

The Data for cell and plasma samples obtained under three experimental conditions (control, vehicle, and treated) were acquired using Agilent MassHunter Data Acquisition software (Version: B.10.1.48) in both ESI modes. The raw data (.d) files were imported to Agilent MassHunter Qualitative Analysis software (Version: B.10.0.1) to assess chromatographic peak shapes, retention time profiles, and mass spectral background noise. The retention time across all data files under each experimental condition for cell and plasma samples under each ESI mode was corrected using internal standard solution analytical runs with a minimum spectral height. This ensured the correction of retention time deviations across different analytical runs and the removal of background noise in the mass spectrum. The batch recursive molecular feature extraction strategy was used to identify molecular features. The specific parameters and values are as follows: the extraction parameters included a minimum mass spectral peak height of 1,500 counts, the allowed ion species of [M+H] +, [M+Na] +, and [M+NH 4] + for the positive ion mode, and [M−H] − for the negative ion mode, the isotope model of common organic molecules without halogens, and the limit assigned charge states to a range of 1-2; the compound filters were set by default; the compound binning and alignment parameters included a retention time tolerance of 0.10% + 0.30 min, and a mass tolerance of 20.00 ppm +2.00 mDa; the post-processing filters were set at an absolute height of at least 5,000 counts for mass spectral peaks, a molecular feature extraction score of at least 75, and a minimum match of molecular feature at 75% (this meant a molecular feature must be present in 3 out of 4 replicate runs in each experimental condition to be included). For the find by ion, the matching tolerance and scoring parameters included a mass score at 100, isotope abundance and spacing scores at 60 and 50, respectively, and a retention score of 0; the EIC peak integration and filtering parameters included an absolute height of at least 7,000 counts for chromatographic peaks; the spectrum extraction and centroiding parameters were set by default; and the post-processing filters included an absolute height of at least 7,000 counts for chromatographic peak heights and a target score of at least 75.00. This was performed by uploading data files of different experimental conditions (control, vehicle, and treated) for cell samples in positive ESI data acquisition mode and separately for negative ESI data acquisition mode. A similar was performed for plasma samples. Finally, the data of all experimental groups by each ionization mode obtained from the operations of “molecular feature extraction” and “find by ion” were exported as profinder archive (.pfa) files from the Agilent MassHunter Profinder software. Multivariate analysis: The reproducibility of the UHPLC-QTOF/MS data was assessed using multivariate principal component analysis (PCA) using MetaboAnalyst V6.0. In detail, using the m/z, retention, and peak area data of all molecular features identified by profinder software different PCA plots were generated. This data was exported in separate .csv files for every experimental condition (i.e., control, vehicle, and treated) in both ESI data acquisition modes (positive and negative) for the samples (cell and plasma). These replicates of the individual condition .csv files were grouped in a folder and zipped together as .zip files (four files, positive ESI mode for cell samples, negative ESI mode for cell samples, Positive ESI mode for plasma samples, and negative ESI mode for plasma samples). These individual .zip files were uploaded to MetaboAnalyst under the statistical section as MS (mass spectrum) peak list data. The retention time and mass tolerance were set to 30.0 seconds and 0.025 Da. The data filtering was performed using an “interquartile range” (IQR) of 5% for removing variables to increase accuracy. Further, the data was normalized using the “internal standard” molecular feature, and the data was log-transformed to the base 10 and auto-scaled. Then, it was submitted for multivariate analysis to generate 2D PCA plots.

#### Metabolite identification and statistical analysis

The (.pfa) files containing molecular feature data were uploaded to Agilent Mass Profiler Professional (MPP) software (Version: B.15.1.2) for statistical analysis and metabolite identification. This was performed separately for positive and negative data acquisition mode for cell samples, and similarly for plasma samples. The metabolomics data uploaded to MPP was normalized using the internal standard (PGE2-d4) spiked in each sample to correct the signal fluctuations between the analytical runs. The data for vehicle and treated cells were adjusted to the median values of the control data to assess the regulation of metabolites under different experimental conditions. The molecular features were annotated using Agilent’s METLIN accurate mass lipids and metabolites libraries with a passing score of 75.0. One-way ANOVA statistical analysis followed by Tukey’s HSD test was applied to identify statistically significant metabolites (*p-value*; ≤ 0.05) with Benjamini-Hochberg correction for FDR. The log2 values of peak areas of the metabolites were used to identify the significantly regulated metabolites between pairs of experimental conditions such as control vs vehicle, control vs. treated, and vehicle vs treated. Metabolites having log2 fold change of greater than 2.0 (or peak area greater than 4 folds) were considered as significantly regulated metabolites between the pair of the noted experimental conditions.

#### A case study on treating a feline leukemia positive/effusive FIP positive cat with molnupiravirrivari and PI

Buddy Boy was a stray domestic short-haired male cat who tested positive for feline leukemia with an IDEXX SNAP combo test during neutering surgery. He was transferred into foster care and developed a distended abdomen within a month. Blood count and chemistry profile were obtained using the ProCyte Dx Hematology Analyzer and the Catalyst One Chemistry Analyzer. The abdominal effusion was subjected to PCR testing for feline coronavirus conducted by IDEXX. Buddy Boy was then placed on oral molnupiravirravir and PI using the protocol summarized in Figure 2.

**Figure 2.**
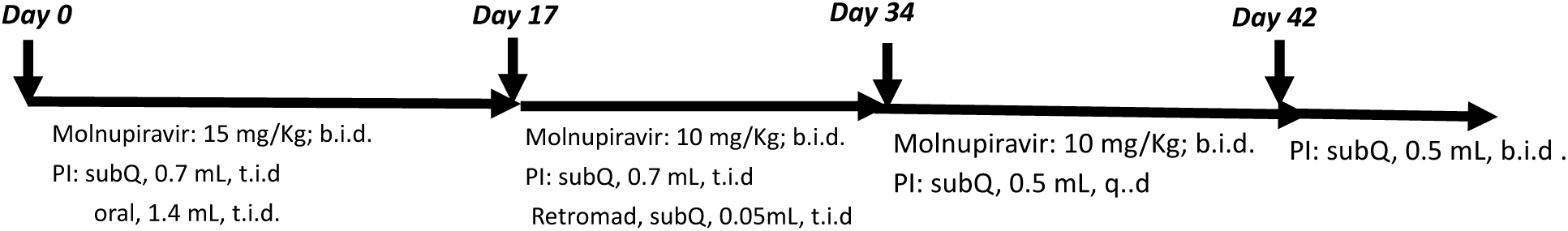

## Results and Discussion

### Metabolomic studies

The biochemical effects of PI were quantified by comparing metabolomic profiles of THP-1 cells treated with PI (ex vivo) with whole blood from mice treated with equivalent concentrations of PI (in vivo). In both ex vivo and in vivo studies, PI-treated samples were compared against n-butanol as the vehicle control. Eight CD-8 positive, healthy adult female mice received 0.5 mg of PI SQ over 7 weeks. All mice were housed in micro-isolators and thus not exposed to or challenged with any pathogens. Thus, the metabolomics profile evaluates the longitudinal effects of PI on non-infected animals. Mice were administered 3mg/kg via IP injection on a Q.D. schedule and were sacrificed 8 hours following the third dose. The respective metabolomic data acquired using an LC-QTOF/MS 6545 on whole blood and THP-1 cells are summarized in Supplementary Figure 1, which shows representative LC-MS chromatograms of the metabolites in the blood of unchallenged mice, mice treated with n-butanol (vehicle control, and PI-treated mice (3 mg/kg). The Agilent MassHunter Profinder software was used for time alignment and signal normalization by exogenous internal standard and molecular feature extraction (batch recursive feature extraction for small molecules). The MPP program was used to achieve internal standard and protein normalization, and statistics analyses, filtered by frequency and background subtraction. Cross-referencing the whole blood and THP-1 cell lysate sample sets using the statistical analysis procedure described in the method section above identified nine common metabolites as summarized in the table shown in Figure 3, with the highest impact score found in the sphingolipid metabolism being 0.3. The impact sore was used to rank the significance of the metabolite match, with the highest value being the most significant. The impact score of sphingolipid metabolism is higher than the other common metabolites identified in whole blood and THP-1 samples; sphinganine and N-acyl sphingosine were the common metabolites upregulated by PI treatment. Sphingolipids are known to be associated with the activation and/or regulation of cell-mediated immunity.^42^ As PI also bears one or two phosphate moieties at the end of its hydrophobic carbon chain like sphingolipids (Figure 1), it possible that PI interacts with some of the protein components in the sphingolipid metabolic pathway are the molecular targets to initiate an immune response. From a statistical standpoint, the impact score of 0.3 is relatively low. But the metabolomic experiments performed in this study were monitored at a specific concentration of PI at only one time. Additional metabolomic experiments conducted at different PI treatment times and concentrations should confirm the participation of PI in sphingolipid metabolism, and reveal the molecular targets.

**Figure 3.**
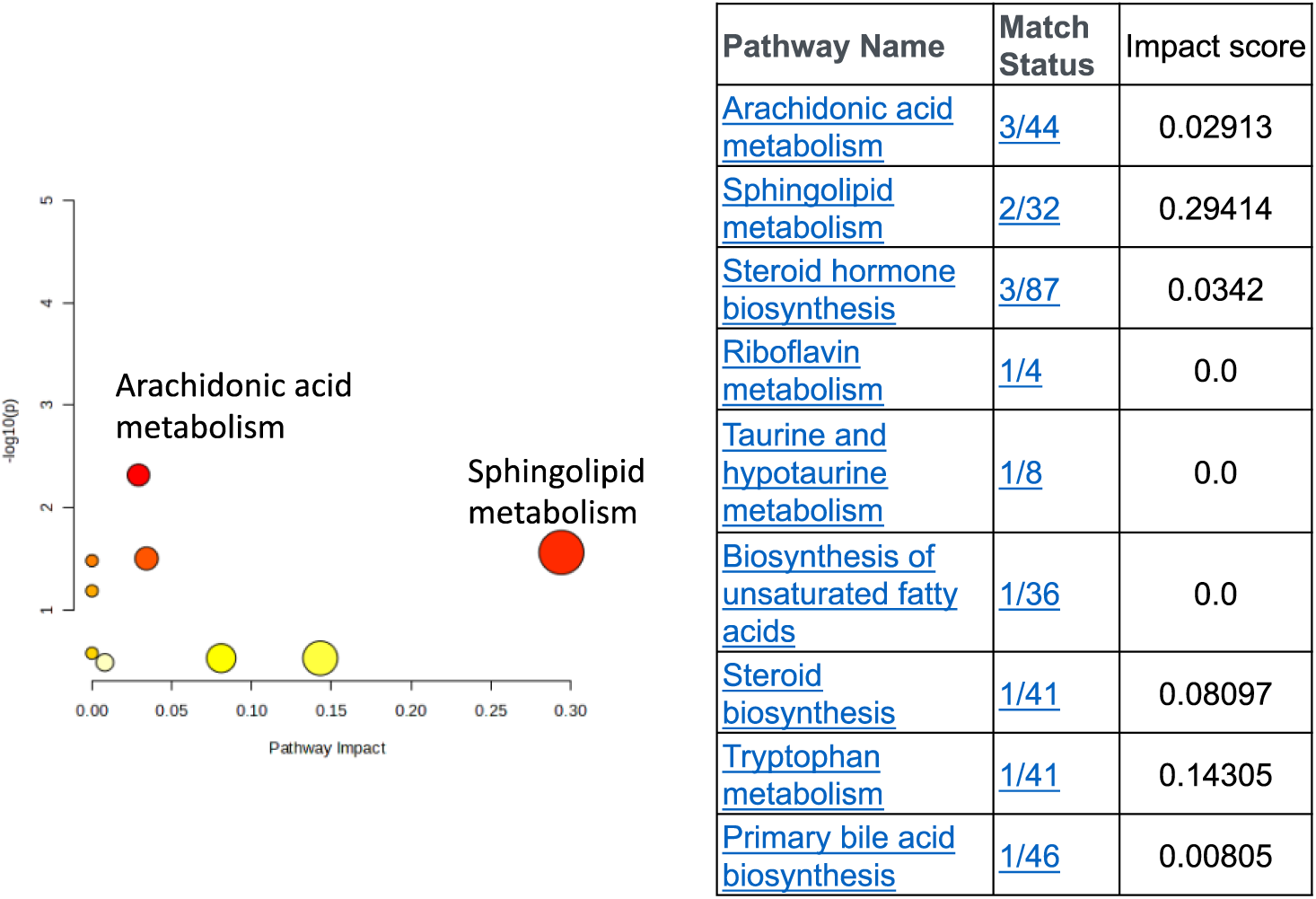

**Figure 4.**
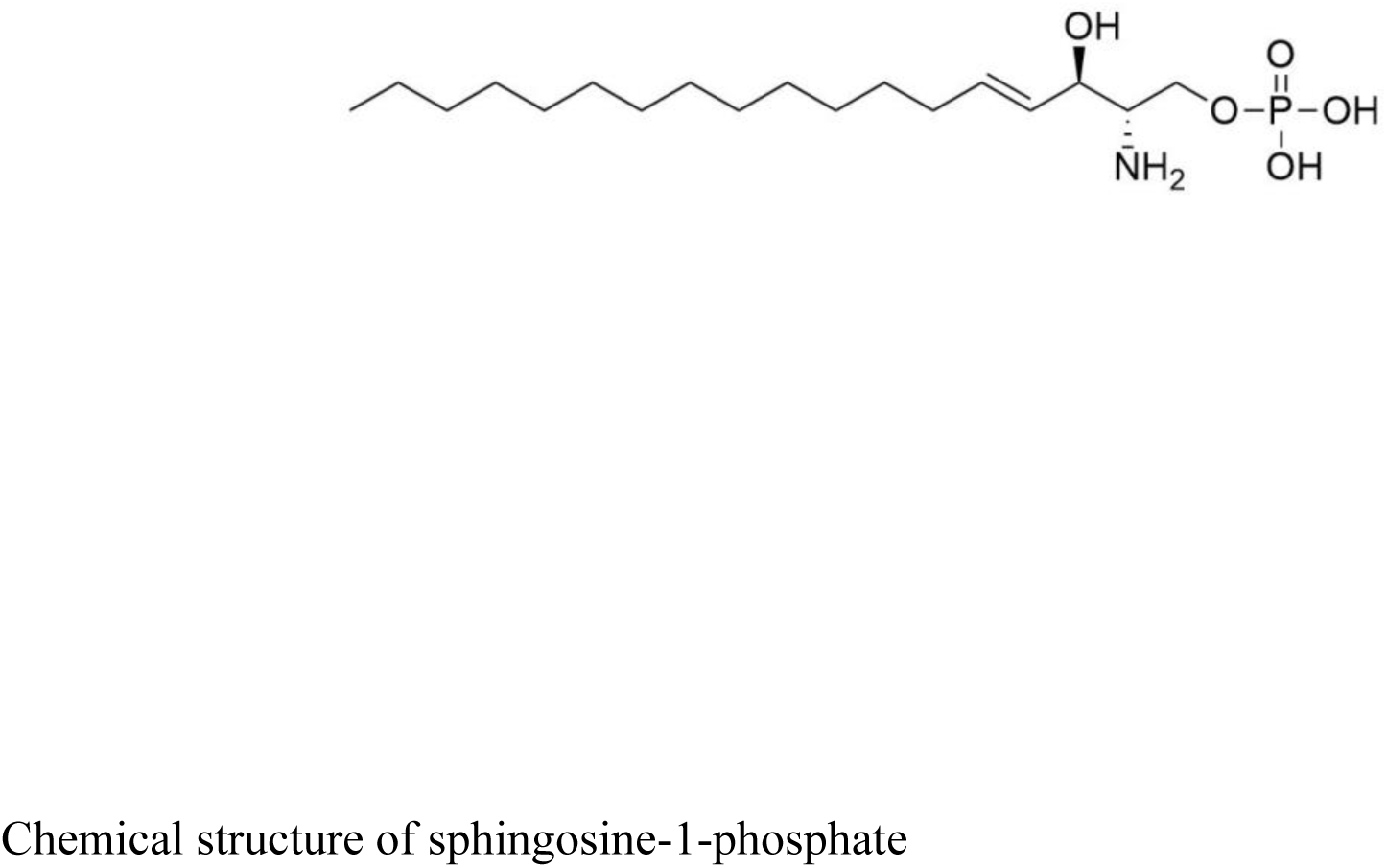
Chemical structure of sphingosine-1-phosphate.

THP-1 cells have been used for decades as a cell model to study inflammation and differentiation into macrophages. The detection of an altered metabolomic profile in the blood of healthy mice treated with PI is interesting. PI has been used as an immunomodulator to combat viral infections in felines. As such, its impact on a healthy animal has not been evaluated until now. The discovery that metabolites associated with sphinganine and N-acyl sphingosine metabolism in PI-treated mice suggest an upregulation of innate immunity, accounting for PI’s immunotherapeutic properties. The use of mice as a translational model will significantly facilitate the identification of the molecular targets of PI, especially in inflammation and age-related disease models. Genetic knock-down and knock-out mice are available for conducting a comparable metabolomic study, and data can be quantified and compared with the findings generated in this study.

### Ex vivo studies examined PI’s biochemical effects against THP-1 cells by flow cytometry

Figure 5 provides a representative dose-response curve examining cellular proliferation as a function of increasing PI concentrations (0-75 ug/mL) over 24 hours. These data show a steady increase in total cell number as a function of increasing PI concentration. The highest dose test (75 ug/mL) showed a 2-fold increase in total cell number. In addition, there was no statistical difference in the percentage of viable (green) versus non-viable cells (black) treated with 75 ug/mL PI versus cells treated with n-butanol (vehicle). This result is consistent with experiments using Annexin V / Propidium Iodide uptake as an early and late-stage apoptosis marker (data not shown).

**Fig. 5.**
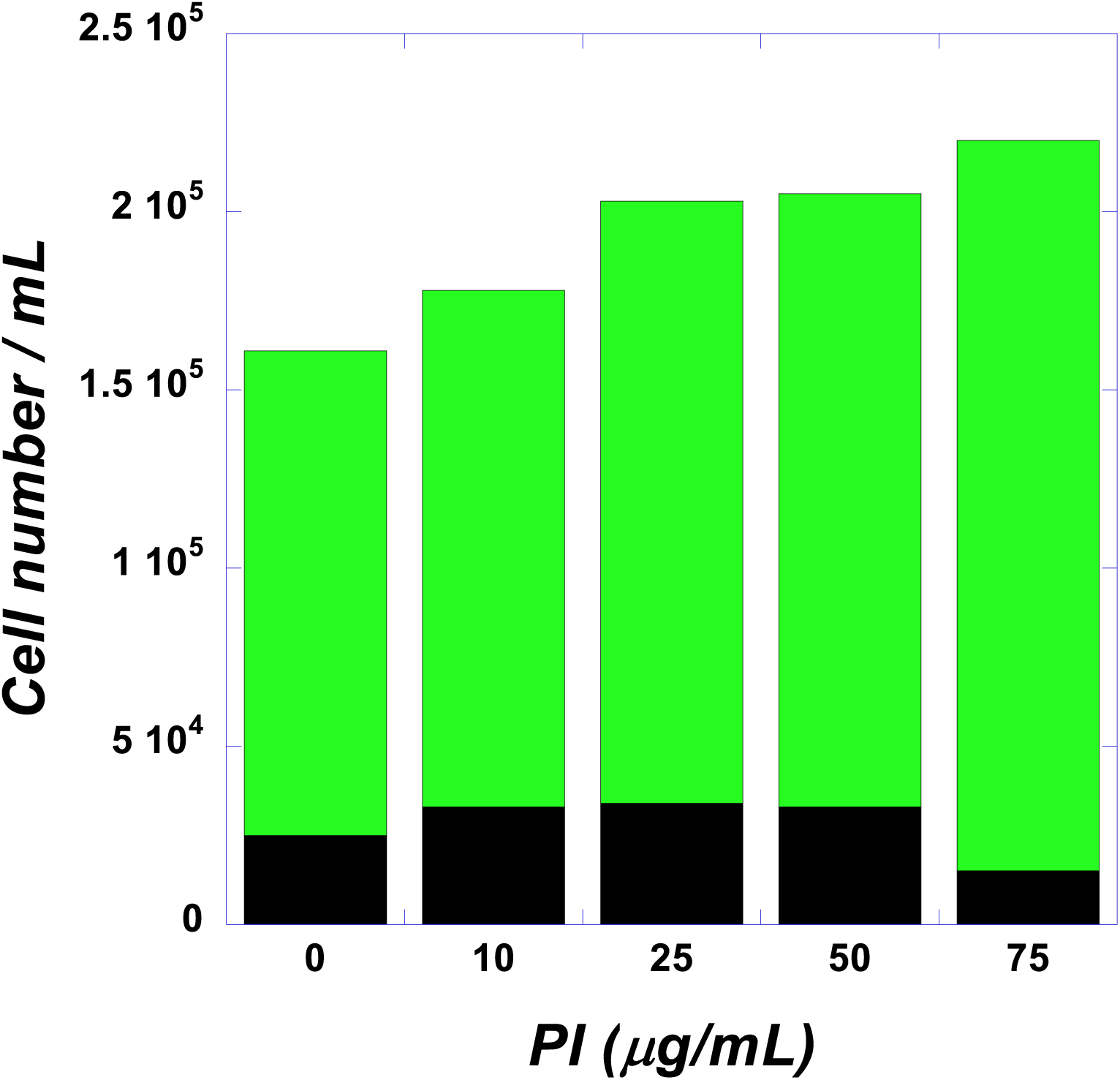
Shows the dose response curve for PI plotting viable (green) and non-viable (black) THP-1 cells as a function of increasing PI concentrations. This was a fixed time point assay taken at 24 hours post treatment.

### Immunophenotypic analyses to quantify whole blood, bone marrow and splenic immune cell populations by flow cytometry

Metabolomic studies of the whole blood samples of healthy mice indicate the upregulation of sphinganine and N-acylsphingoine metabolisms, both of which contribute to the regulation of the T-cell mediated innate immunity. To corroborate, immunophenotyping flow cytometry analyses were performed in the whole blood samples harvested from the same mice used in the metabolomic studies. Each blood sample was stained with the myeloid/lymphoid antibody markers and detected by fluorescent flow cytometry. Figure 6 summarizes the flow cytometry data for the same mice blood samples used in the metabolomic studies. Overall, the red blood cell and B-cell counts remain relatively constant in all the samples. However, from the CBC data, there is a trend toward increased total white cells, neutrophils, and lymphocytes in PI versus placebo mice (Figure 6A), accompanied by increased myeloid cells, decreased lymphoid cells, and decreased total T cells (Figure 6B).

**Figure 6.**
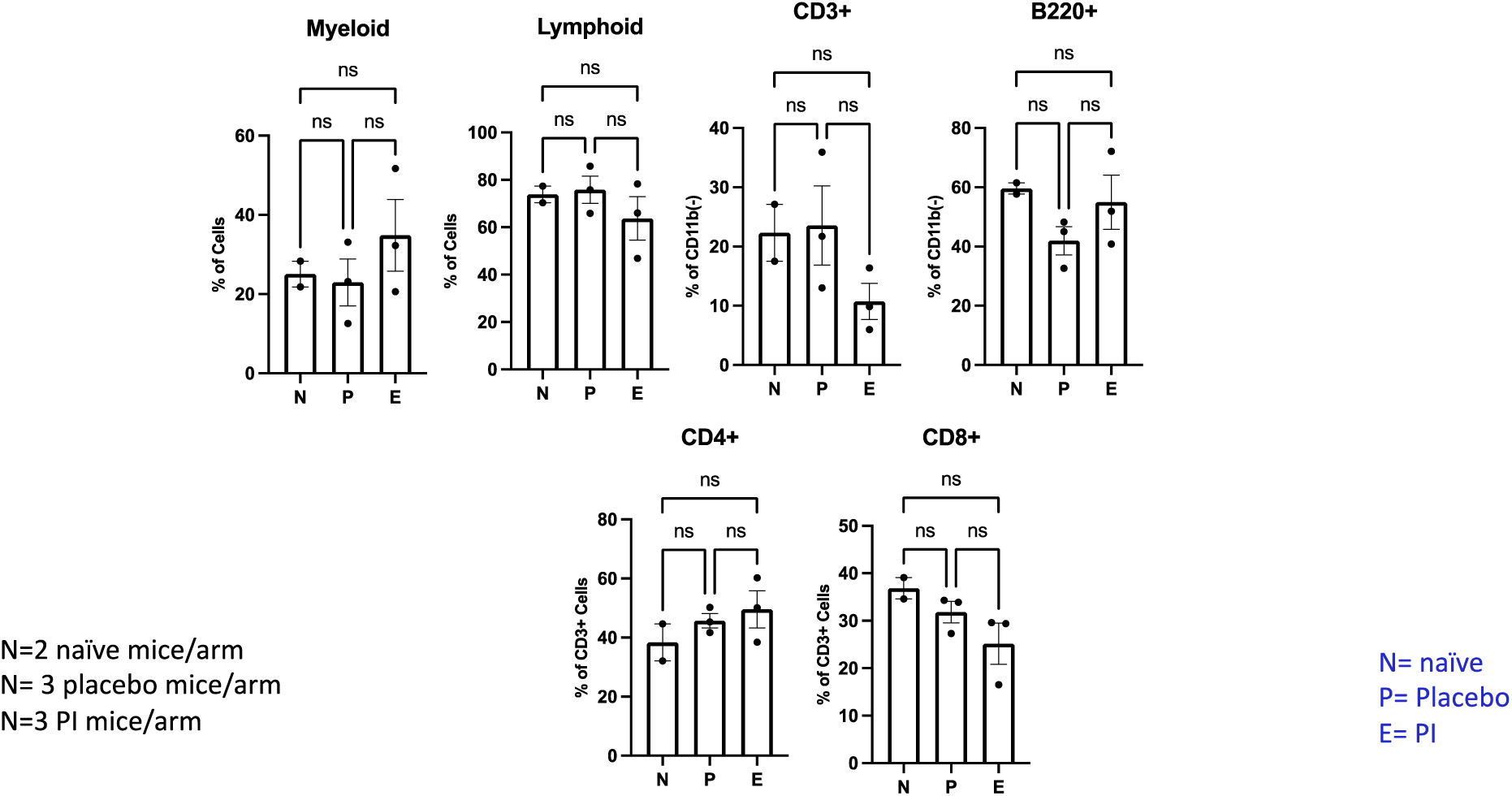
Polyprenyl immunostimulant (PI) moderately impacts the frequencies of B and T cells in peripheral blood.

There are increased CD4+ and decreased CD8+ cells within the T cell populations, hence an overall increase in CD4/CD8 ratio in the cell profile. In human immunodeficiency, an increased CD4/CD8 ratio in CBC is considered an improvement in immunity.^43^ The detection of an increase in CD4/CD8 in the PI-condition healthy mice blood further the immunotherapeutic potential of this compound. However, it should be noted that this study did not include a sampling size to determine statistical relevance. Nevertheless, when interpreted along with the metabolomic data and the steady-state hematopoiesis study described below, this result implicates the impact of PI on T-cell-mediated immunity.

To expand the sampling size and subject diversity, ten 9-weeks old female C57/Bl6 mice that had never been exposed to PI were treated with the same protocol as the aged mice used in the metabolomic study. Peripheral blood, bone marrow, and splenic tissue were harvested, and cells were stained with fluorescently labeled antibodies for immunomarkers, followed by flow cytometry imaging. Figure 7 summarizes the flow cytometry results. Figure 7A shows that the cellular composition in the peripheral blood between PI and n-butanol is comparable, indicating that this parameter is not affected. Figure 7B shows the flow cytometry results of the bone marrow (BM) hematopoietic stem and progenitor cells (HSPCs); PI does not significantly impact the cell populations. Figure 7C shows that PI decreases BM B-cells and BM CD8+ cells, but increases BM CD4+ cells, thereby increseing the CD4/CD8 ration. Figure 7D shows that that PI does not affect the population of splenic HSPCs. Figure 7E shows that PI decreases splenic CD8+ cells but does not alter the splenic CD4+ cells, resulting in an overall increase in the CD4/CD8 ratio. In the HIV field, the CD4/CD8 ratio serves as a biomarker to track the robustness of the host’s immune function, with an increase in the value signifying immune activation and immunosenescence.^44^ The detection of an increase in the CD4/CD8 ratio indicates that PI functions to heighten the immune response even in an uninfected animal, which is a valuable property to exploit as an immunotherapeutic agent to combat infectious diseases.

**Figure 7A.**
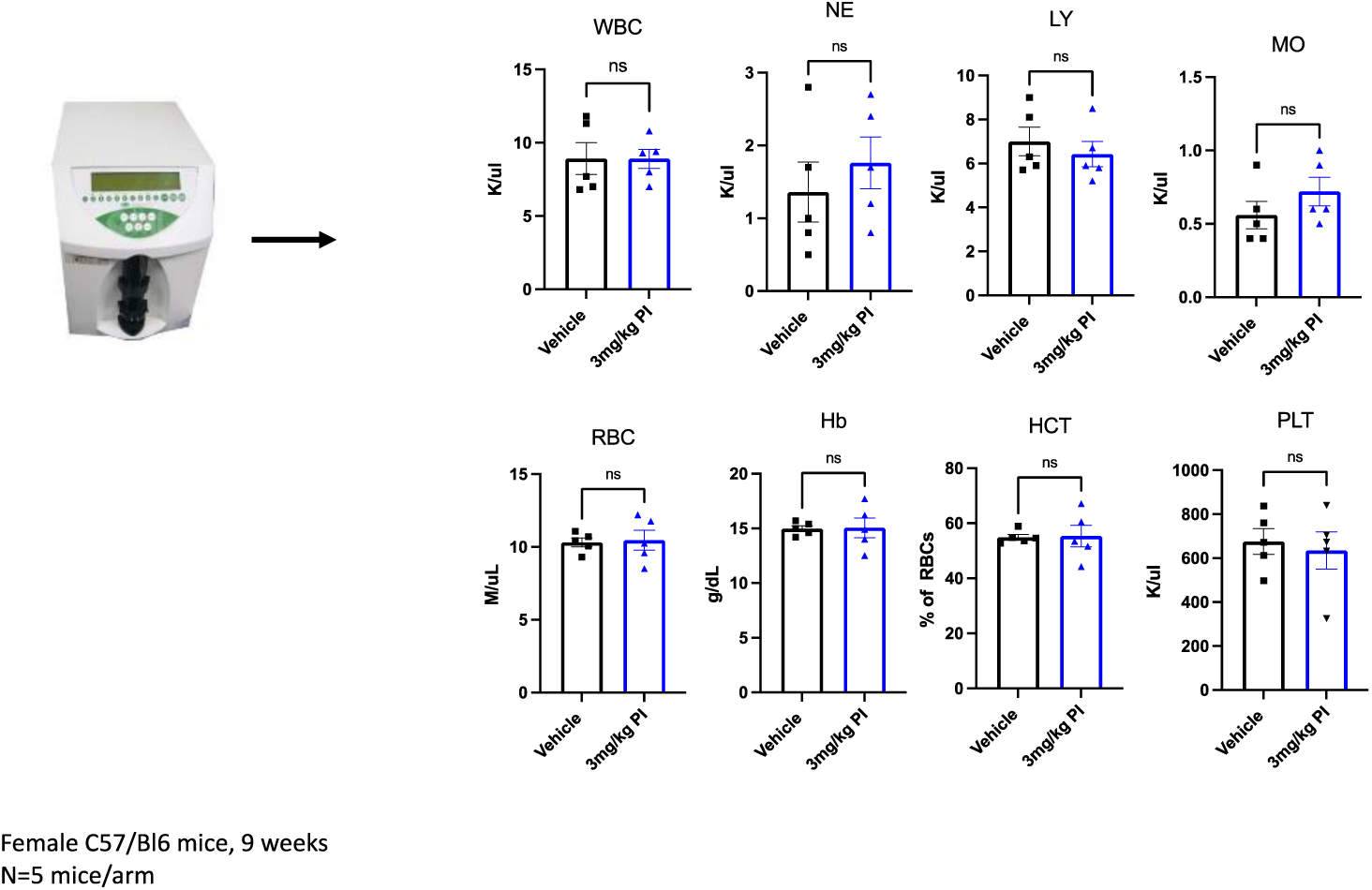
Polyprenyl immunostimulant (PI) does not impact the composition of peripheral blood.

**Figure 7B.**
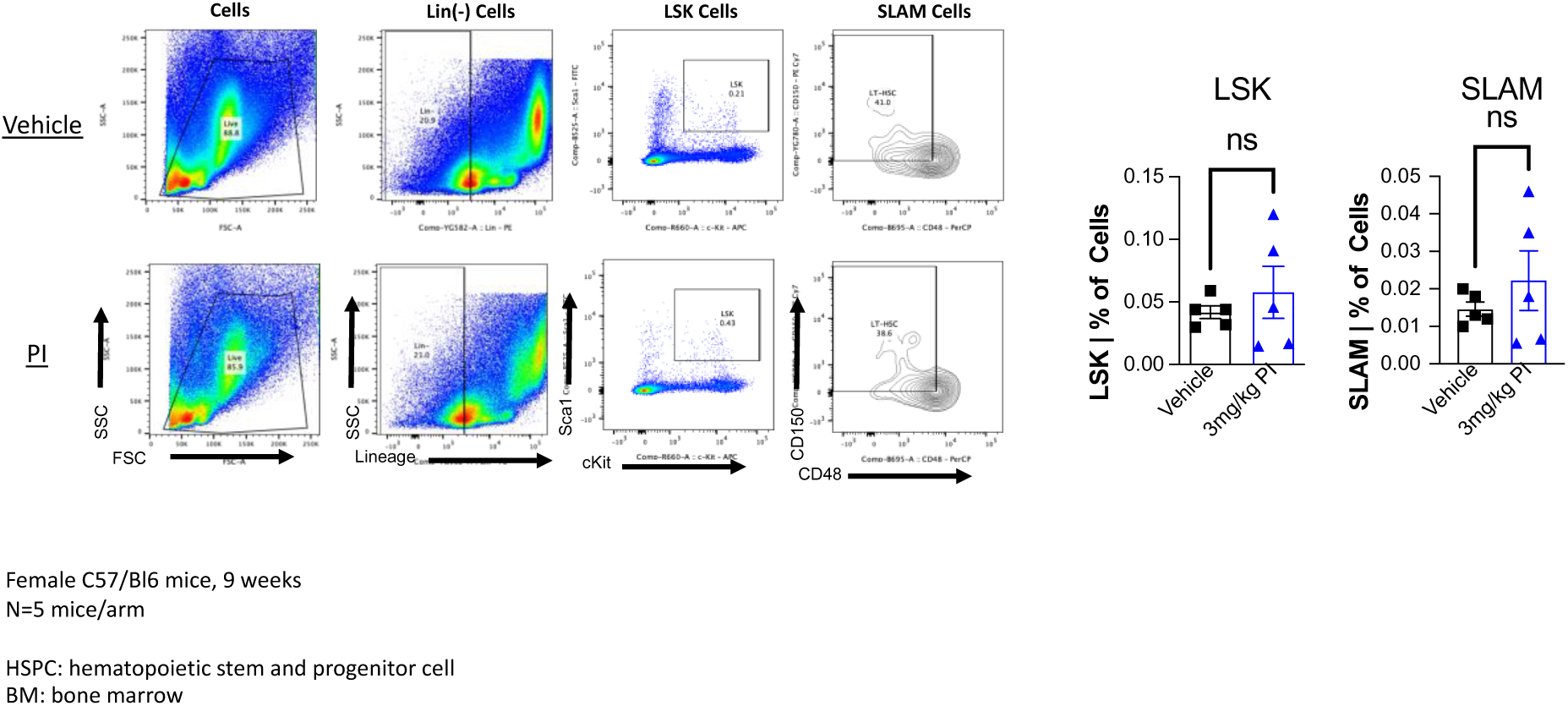
Polyprenyl immunostimulant (PI) does not significantly impact BM HSPC frequencies.

**Figure 7C.**
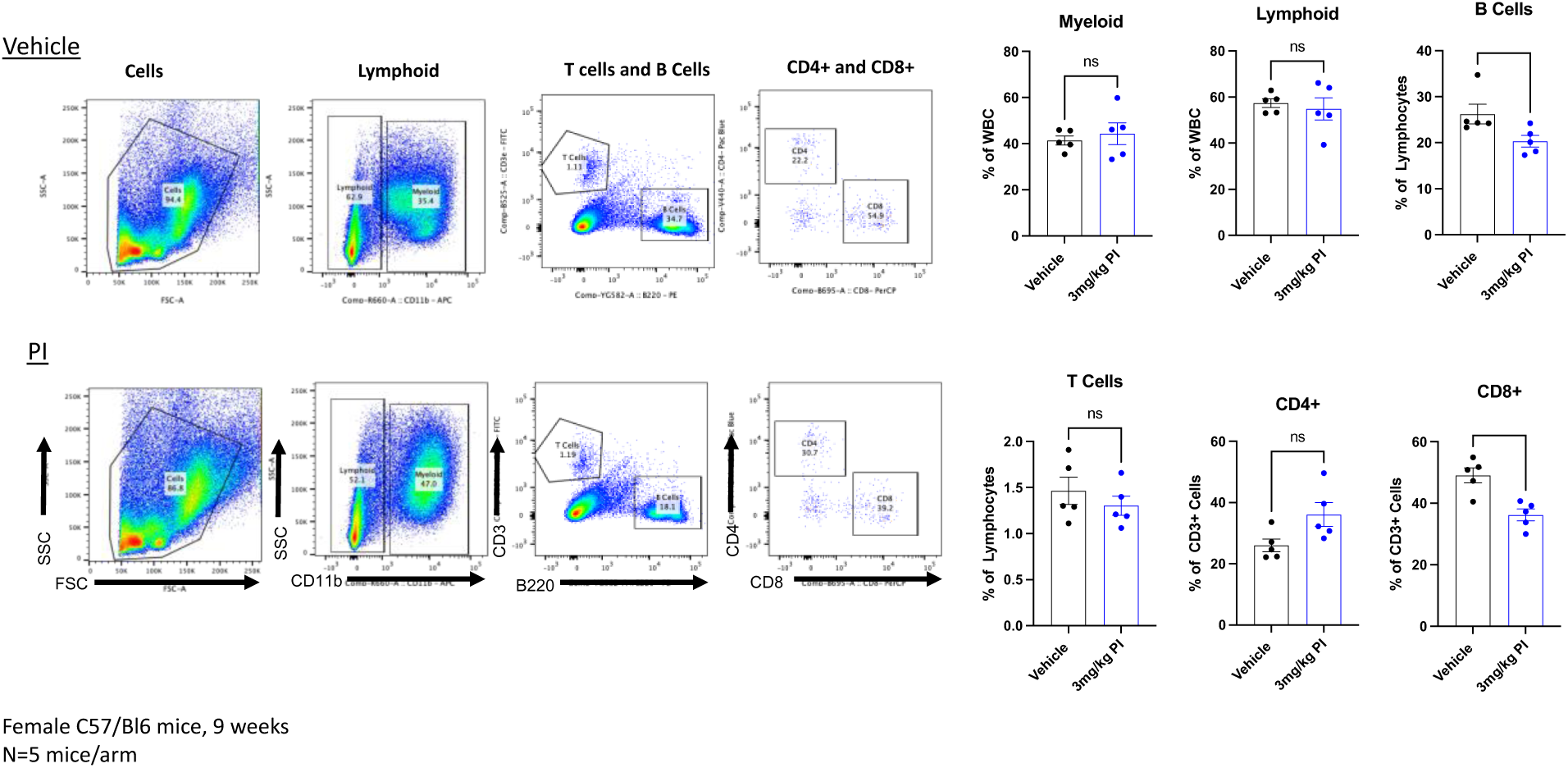
Polyprenyl immunostimulant (PI) reduces BM B-cells and alters BM T-cel frequencies.

**Figure 7D.**
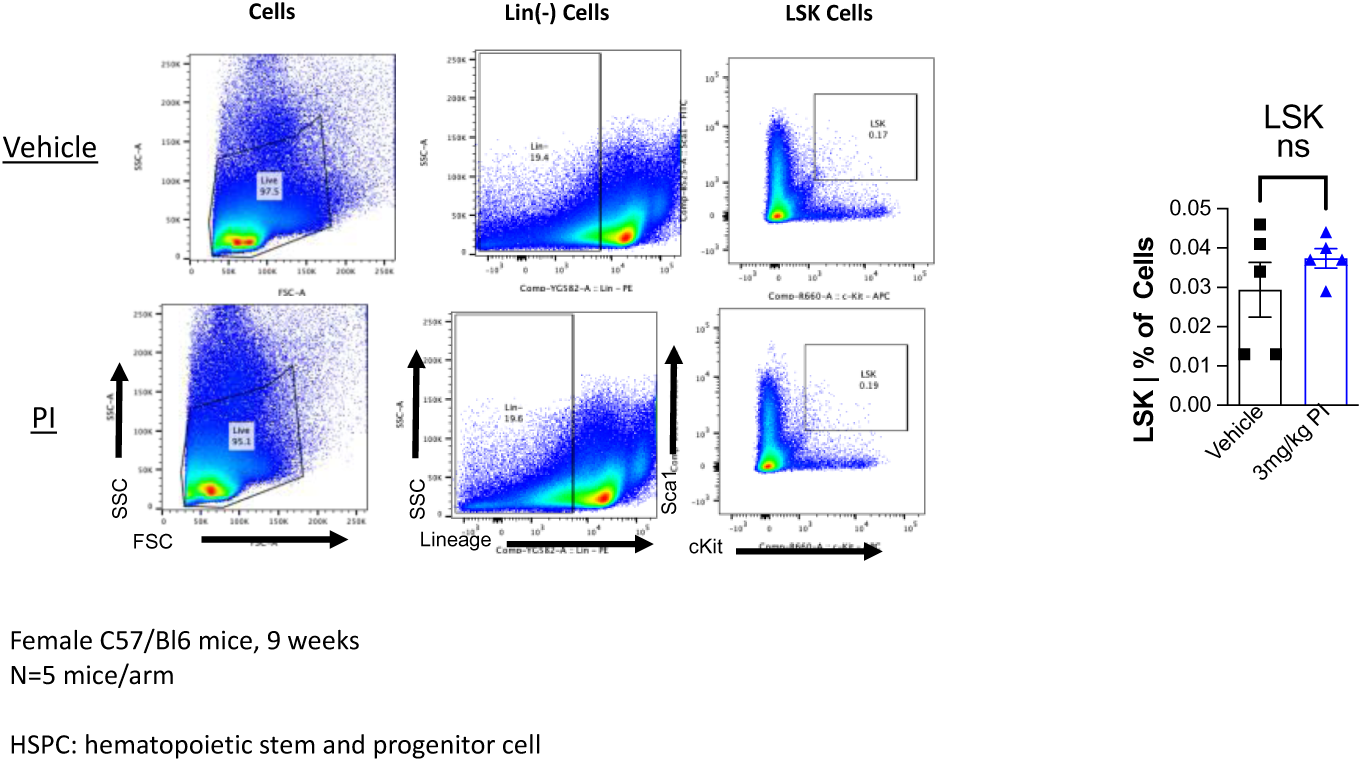
Polyprenyl immunostimulant (PI) does not significantly impact splenic HSPC frequencies.

**Figure 7E.**
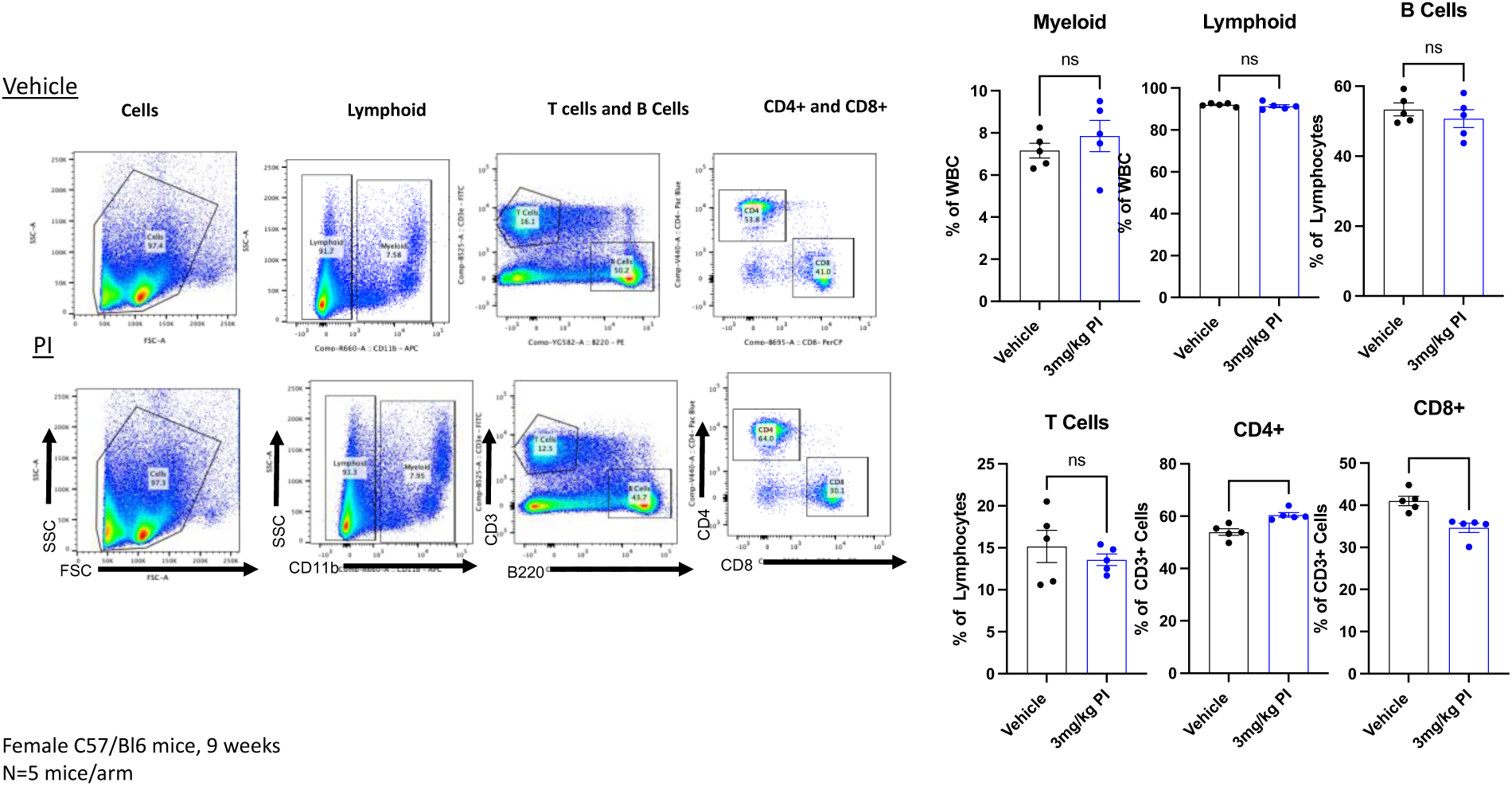
Polyprenyl immunostimulant (PI) alters splenic T-cell frequencies.

### A case study on using PI to enhance the therapeutic property of Molnupiravirrivir

In light of the whole blood and THP-1 cell studies, along with published data showing that PI modulates the innate immunity of an animal,^45^ we conducted a proof of principle study to evaluate the ability of PI to enhance the therapeutic potential of the antiviral drug molnupiravirrivavri (molnupiravir). In humans, molnupiravir showed efficacy in alleviating mild symptoms of COVID. In felines, molnupiravir has been used successfully as an oral medication to treat FIP in non-immune compromised hosts.^33, 46, 47^ However, no study has been found on the efficacy of molnupiravir in treating immune-compromised hosts. For a proof-of-principle study, we treated a stray feline displaying the effusive form of FIP and was confirmed positive for the progressive stage of feline leukemia viral infection with a combination of PI and molnupiravir. Since FIP is fatal when left untreated, we did not include a control for comparison for ethical reasons. Furthermore, the subject of interest here was found to be a 2-year-old stray with an unknown history. Finding a comparable feline to serve as a control would be impossible. To assess the effect of PI, we compared the duration needed to bring the subject feline to remission with the published duration of using molnupiravir to treat unimmune-compromised cats.^33, 46, 47^ Using the protocol outlined in Figure 2, we monitored the FeLV status by the IDEXX FeLv Quant real PCR, the FIP status, and the progression of both diseases by the CBC and chemistry profile. The IDEXX FeLV Quant real PCR test value obtained on day 17 and day 31 of the molnupiravir/PI combo treatment was 14.19 x 10^6^ copies/mL and 41.97 x 10^6^ copies/mL, respectively (data not shown). The quant real PCR value for FeLV did not change significantly during the course of treatment. Table 1 shows the blood count and chemistry profile values that change during the treatment duration. There was a drop in the HCT value after the administration of PI/molnupiravir. Non-regenerative anemia is often a phenotype of FeLv or FIP infection. Since the subject was already receiving PI and molnupiravir to treat FIP, we attributed the drop in HCT to a manifestation of FeLV infection exacerbated by FIP. As such, we added a low dosage of the antiviral agent RetroMad 1 to control the FeLV flareup.^48^ After 5 weeks of treatment, the CBC and chemistry profiles of the subject returned to normal ranges. The abdominal exudate caused by the effusive form of FIP was eliminated, but the progressive stage of FeLv infection persisted. The five-week treatment time needed to attain remission in FIP is less than the reported 12-week protocol for non-FeLV subjects.

**Table 1.** Comparing the blood work parameters of Buddy Boy during FIP treatment.

|  | ← during treatment → |  |  | post-treatment |  |
| --- | --- | --- | --- | --- | --- |
|  | Day 0 | Day 17 | Day 31 | Day 56 | normal range |
| Hematocrit (%) | 26.7 | 22.5 | 33.6 | 41.2 | 30.3-52.3 |
| Neutrophils (K/μL) | 10.61 | 6.12 | 9.37 | 3.65 | 2.3-10.29 |
| Eosinophils (K/μL) | 0 | 0.17 | 0.63 | 0.22 | 0.17-1.57 |
| BUN (mg/dL) | 12 | 20 | 21 | 23 | 16-36 |
| total protein (g/dL) | 7.0 | 9.5 | 7.7 | 7.2 | 5.7-8.9 |
| Globulin (g/dL) | 4.8 | 6.4 | 5.0 | 3.9 | 2.8-5.1 |
| ALP (U/L) | <10 | 54 | 43 | 103 | 14-111 |
| total bilirubin | 1.5 | 0.7 | 0.3 | < 0.1 | 0-0.9 |

Data were generated by the ProCyte Dx Hematology Analyzer and the Catalyst One Chemistry Analyzer abnormal values are highlighted in red

### Conclusion

In this study, we conducted a series of proof-of-principality studies to identify biochemical and/or biomarkers that can be used to track the performance of PI in relation to immune functions. To this end, we utilized metabolomics to identify sphinganine and acylsphingosine, which were upregulated by PI in healthy mice and in THP-1 cells, with both metabolites found in the sphingolipid metabolic pathway, which plays a significant role in innate immunity in mammals. Immunophenotypic analyses by fluorescent flow cytometry were used to corroborate the PI-upregulated pathways. Since the mice and THP-1 cells used in this study were examined under insult-free conditions, the quantitative nature of metabolomic and flow cytometry allows the setting of reference standards in the insult-free samples such that the effect of PI in combating viral infection or other hematopoietic injury can be quantitatively accessed. A proof-of-principle case study was performed to demonstrate that adding PI to the molnupiravir therapy reduces the treatment duration of an immunopromised subject. Collectively, the findings generated from this study establish the premise and the approaches for studying the molecular mechanism of PI and evaluate the ability of PI to synergize the therapeutic efficacies of other antiviral drugs in general.

## Author Contributions

Conceptualization, Lee, I, Berdis, A.J., Desai, A and Xu, Y; methodology, Berdis, A.J., Patil, A and Desai, A., Pozza-Adams, K.; data analysis, Berdis, A.J., Patil, A and Desai, A., Pozza-Adams, K, Xu, Y. and Lee, I.; writing-original draft preparation, Lee, I.; writing-review and editing, Berdis, A.J., Patil, A and Desai, A., Pozza-Adams, K, Xu, Y. and Lee, I. All authors have read and agreed to the published version of the manuscript.

## Acknowledgments

We wish to thank Sass&Sass, Inc. and Gina Galyon for the donation of CD-1 mice blood serum, and Ms Fabienne Ridolfi for assistance in establishing a protocol for treating Buddy Boy for FIP.

Institutional Review Board Statement

Experimental procedures were approved by the CWRU Institutional Animal Care and Use Committee (IACUC) in accordance with approved IACUC protocols (2019-0065).

## References

1. Banerjee A, Kulcsar K, Misra V, Frieman M and Mossman K. Bats and Coronaviruses. Viruses. 2019;11.

2. Cui J, Li F and Shi ZL. Origin and evolution of pathogenic coronaviruses. Nat Rev Microbiol. 2019;17:181–192.

3. Fehr AR and Perlman S. Coronaviruses: an overview of their replication and pathogenesis. Methods Mol Biol. 2015;1282:1–23.

4. Shi Z and Hu Z. A review of studies on animal reservoirs of the SARS coronavirus. Virus Res. 2008;133:74–87.

5. Masters PS. The molecular biology of coronaviruses. Adv Virus Res. 2006;66:193–292.

6. Weiss SR and Leibowitz JL. Coronavirus pathogenesis. Adv Virus Res. 2011;81:85–164.

7. Abdul-Rasool S and Fielding BC. Understanding Human Coronavirus HCoV-NL63. Open Virol J. 2010;4:76–84.

8. Corman VM, Muth D, Niemeyer D and Drosten C. Hosts and Sources of Endemic Human Coronaviruses. Adv Virus Res. 2018;100:163–188.

9. Lau SK, Lee P, Tsang AK, Yip CC, Tse H, Lee RA, So LY, Lau YL, Chan KH, Woo PC and Yuen KY. Molecular epidemiology of human coronavirus OC43 reveals evolution of different genotypes over time and recent emergence of a novel genotype due to natural recombination. J Virol. 2011;85:11325–37.

10. Lau SK, Woo PC, Yip CC, Tse H, Tsoi HW, Cheng VC, Lee P, Tang BS, Cheung CH, Lee RA, So LY, Lau YL, Chan KH and Yuen KY. Coronavirus HKU1 and other coronavirus infections in Hong Kong. J Clin Microbiol. 2006;44:2063–71.

11. Gaunt ER, Hardie A, Claas EC, Simmonds P and Templeton KE. Epidemiology and clinical presentations of the four human coronaviruses 229E, HKU1, NL63, and OC43 detected over 3 years using a novel multiplex real-time PCR method. J Clin Microbiol. 2010;48:2940-7.

12. Chafekar A and Fielding BC. MERS-CoV: Understanding the Latest Human Coronavirus Threat. Viruses. 2018;10.

13. Cheng VC, Chan JF, To KK and Yuen KY. Clinical management and infection control of SARS: lessons learned. Antiviral Res. 2013;100:407–19.

14. Desforges M, Le Coupanec A, Dubeau P, Bourgouin A, Lajoie L, Dube M and Talbot PJ. Human Coronaviruses and Other Respiratory Viruses: Underestimated Opportunistic Pathogens of the Central Nervous System? Viruses. 2019;12.

15. Mackay IM and Arden KE. MERS coronavirus: diagnostics, epidemiology and transmission. Virol J. 2015;12:222.

16. Momattin H, Mohammed K, Zumla A, Memish ZA and Al-Tawfiq JA. Therapeutic options for Middle East respiratory syndrome coronavirus (MERS-CoV)--possible lessons from a systematic review of SARS-CoV therapy. Int J Infect Dis. 2013;17:e792–8.

17. Satija N and Lal SK. The molecular biology of SARS coronavirus. Ann N Y Acad Sci. 2007;1102:26–38.

18. Song Z, Xu Y, Bao L, Zhang L, Yu P, Qu Y, Zhu H, Zhao W, Han Y and Qin C. From SARS to MERS, Thrusting Coronaviruses into the Spotlight. Viruses. 2019;11.

19. Weiss SR and Navas-Martin S. Coronavirus pathogenesis and the emerging pathogen severe acute respiratory syndrome coronavirus. Microbiol Mol Biol Rev. 2005;69:635–64.

20. Ahn DG, Shin HJ, Kim MH, Lee S, Kim HS, Myoung J, Kim BT and Kim SJ. Current Status of Epidemiology, Diagnosis, Therapeutics, and Vaccines for Novel Coronavirus Disease 2019 (COVID-19). J Microbiol Biotechnol. 2020;30:313-324.

21. Guo YR, Cao QD, Hong ZS, Tan YY, Chen SD, Jin HJ, Tan KS, Wang DY and Yan Y. The origin, transmission and clinical therapies on coronavirus disease 2019 (COVID-19) outbreak - an update on the status. Mil Med Res. 2020;7:11.

22. Tian S, Hu W, Niu L, Liu H, Xu H and Xiao SY. Pulmonary Pathology of Early-Phase 2019 Novel Coronavirus (COVID-19) Pneumonia in Two Patients With Lung Cancer. J Thorac Oncol. 2020;15:700–704.

23. Jackwood MW. The relationship of severe acute respiratory syndrome coronavirus with avian and other coronaviruses. Avian Dis. 2006;50:315–20.

24. Hodgson T, Britton P and Cavanagh D. Neither the RNA nor the proteins of open reading frames 3a and 3b of the coronavirus infectious bronchitis virus are essential for replication. J Virol. 2006;80:296–305.

25. Barr F. Feline infectious peritonitis. J Small Anim Pract. 1998;39:501–4.

26. Kipar A and Meli ML. Feline infectious peritonitis: still an enigma? Vet Pathol. 2014;51:505–26.

27. Pedersen NC. An update on feline infectious peritonitis: diagnostics and therapeutics. Vet J. 2014;201:133–41.

28. McGonagle D, Sharif K, O’Regan A and Bridgewood C. The Role of Cytokines including Interleukin-6 in COVID-19 induced Pneumonia and Macrophage Activation Syndrome-Like Disease. Autoimmun Rev. 2020;19:102537.

29. Soy M, Keser G, Atagunduz P, Tabak F, Atagunduz I and Kayhan S. Cytokine storm in COVID-19: pathogenesis and overview of anti-inflammatory agents used in treatment. Clin Rheumatol. 2020;39:2085–2094.

30. Ye Q, Wang B and Mao J. The pathogenesis and treatment of the; Cytokine Storm’ in COVID-19. J Infect. 2020;80:607–613.

31. Beigel JH, Tomashek KM, Dodd LE, Mehta AK, Zingman BS, Kalil AC, Hohmann E, Chu HY, Luetkemeyer A, Kline S, Lopez de Castilla D, Finberg RW, Dierberg K, Tapson V, Hsieh L, Patterson TF, Paredes R, Sweeney DA, Short WR, Touloumi G, Lye DC, Ohmagari N, Oh MD, Ruiz-Palacios GM, Benfield T, Fatkenheuer G, Kortepeter MG, Atmar RL, Creech CB, Lundgren J, Babiker AG, Pett S, Neaton JD, Burgess TH, Bonnett T, Green M, Makowski M, Osinusi A, Nayak S, Lane HC and Members A-SG. Remdesivir for the Treatment of Covid-19 - Final Report. N Engl J Med. 2020;383:1813–1826.

32. Tian L, Pang Z, Li M, Lou F, An X, Zhu S, Song L, Tong Y, Fan H and Fan J. Molnupiravir and Its Antiviral Activity Against COVID-19. Front Immunol. 2022;13:855496.

33. Reagan KL, Brostoff T, Pires J, Rose A, Castillo D and Murphy BG. Open label clinical trial of orally administered molnupiravir as a first-line treatment for naturally occurring effusive feline infectious peritonitis. J Vet Intern Med. 2024;38:3087–3094.

34. Legendre AM, Kuritz T, Heidel RE and Baylor VM. Polyprenyl Immunostimulant in Feline Rhinotracheitis: Randomized Placebo-Controlled Experimental and Field Safety Studies. Front Vet Sci. 2017;4:24.

35. Cerna P, Ayoob A, Baylor C, Champagne E, Hazanow S, Heidel RE, Wirth K, Legendre AM and Gunn-Moore DA. Retrospective Survival Analysis of Cats with Feline Infectious Peritonitis Treated with Polyprenyl Immunostimulant That Survived over 365 Days. Pathogens. 2022;11.

36. Legendre AM and Bartges JW. Effect of Polyprenyl Immunostimulant on the survival times of three cats with the dry form of feline infectious peritonitis. J Feline Med Surg. 2009;11:624–6.

37. Legendre AM, Kuritz T, Galyon G, Baylor VM and Heidel RE. Polyprenyl Immunostimulant Treatment of Cats with Presumptive Non-Effusive Feline Infectious Peritonitis In a Field Study. Front Vet Sci. 2017;4:7.

38. Sykes JE and Hartmann K. Feline Leukemia Virus Infection. Canine and Feline Infectious Diseases. 2014:224–38.

39. Tasker S, Addie DD, Egberink H, Hofmann-Lehmann R, Hosie MJ, Truyen U, Belák S, Boucraut-Baralon C, Frymus T, Lloret A, Marsilio F, Pennisi MG, Thiry E, Möstl K and Hartmann K. Feline Infectious Peritonitis: European Advisory Board on Cat Diseases Guidelines. Viruses. 2023;15.

40. Kuritz T. Methods and compositions for modulation of innate immunity. 2018;US9872867B2.

41. Paltrinieri S, Giordano A, Stranieri A and Lauzi S. Feline infectious peritonitis (FIP) and coronavirus disease 19 (COVID-19): Are they similar? Transbound Emerg Dis. 2021;68:1786–1799.

42. Lee M, Lee SY and Bae Y-S. Functional roles of sphingolipids in immunity and their implication in disease. Experimental & Molecular Medicine. 2023;55:1110–1130.

43. Bruno G, Saracino A, Monno L and Angarano G. The Revival of an “Old” Marker: CD4/CD8 Ratio. AIDS Rev. 2017;19:81–88.

44. Lu W, Mehraj V, Vyboh K, Cao W, Li T and Routy JP. CD4:CD8 ratio as a frontier marker for clinical outcome, immune dysfunction and viral reservoir size in virologically suppressed HIV-positive patients. J Int AIDS Soc. 2015;18:20052.

45. Cerna P, Dow S, Wheat W, Chow L, Hawley J and Lappin MR. Comparison of Antiviral Immune Responses in Healthy Cats Induced by Two Immune Therapeutics. Pathogens. 2024;13.

46. Roy M, Jacque N, Novicoff W, Li E, Negash R and Evans SJM. Unlicensed Molnupiravir is an Effective Rescue Treatment Following Failure of Unlicensed GS-441524-like Therapy for Cats with Suspected Feline Infectious Peritonitis. Pathogens. 2022;11.

47. Zuzzi-Krebitz AM, Buchta K, Bergmann M, Krentz D, Zwicklbauer K, Dorsch R, Wess G, Fischer A, Matiasek K, Honl A, Fiedler S, Kolberg L, Hofmann-Lehmann R, Meli ML, Spiri AM, Helfer-Hungerbuehler AK, Felten S, Zablotski Y, Alberer M, Both UV and Hartmann K. Short Treatment of 42 Days with Oral GS-441524 Results in Equal Efficacy as the Recommended 84-Day Treatment in Cats Suffering from Feline Infectious Peritonitis with Effusion-A Prospective Randomized Controlled Study. Viruses. 2024;16.

48. Ung H, Silva P and Floriano A. Case studies on RetroMAD1™, an orally administered recombinant chimeric protein to treat naturally infected Feline Leukemia Virus (FeLV) cats. Archives of Veterinary Science and Medicine. 2019;02.

