## Supplementary Figures for "Pre-Clinical Evaluation of a Novel Immunomodulator: a potential immunotherapy for coronaviral disease"

**Supplementary Figure**: Chromatograms of the metabolites detected by LC-QTOF/MS 6545 in THP-1 cells.

Metabolites recovered in the aqueous layer of the THP-1 cell extraction process under negative ion detection mode. Control: untreated cell; PI: cells treated with PI

Neg_Aq

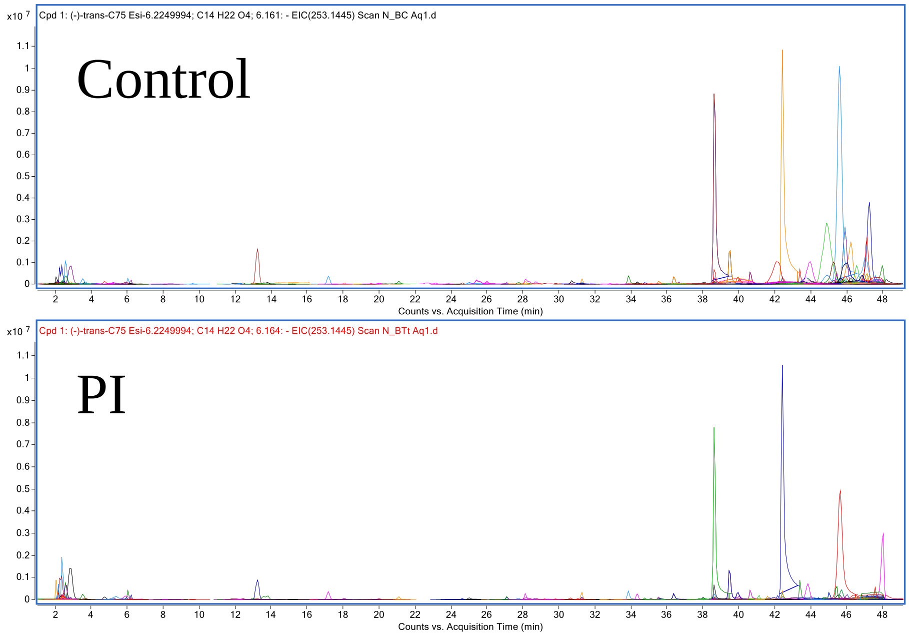

Metabolites recovered in the aqueous layer of the THP-1 cell extraction process under negative ion detection mode. Butanol: cells treated with n-butanol, the vehicle; PI: cells treated with PI

Neg_Aq

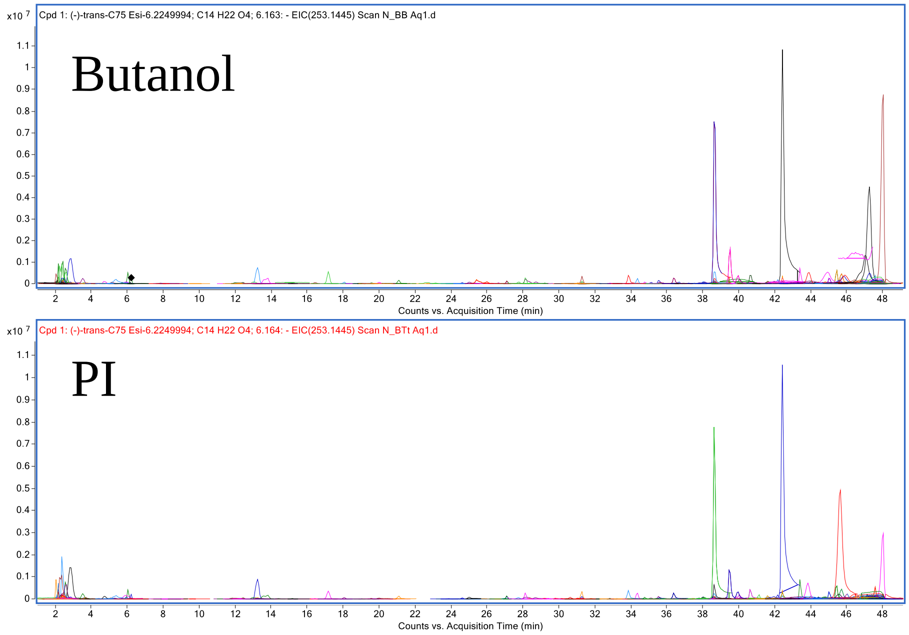

Metabolites recovered in the organic layer of the THP-1 cell extraction process under negative ion detection mode. Control: untreated cells, PI: cells treated with PI

Neg_Org

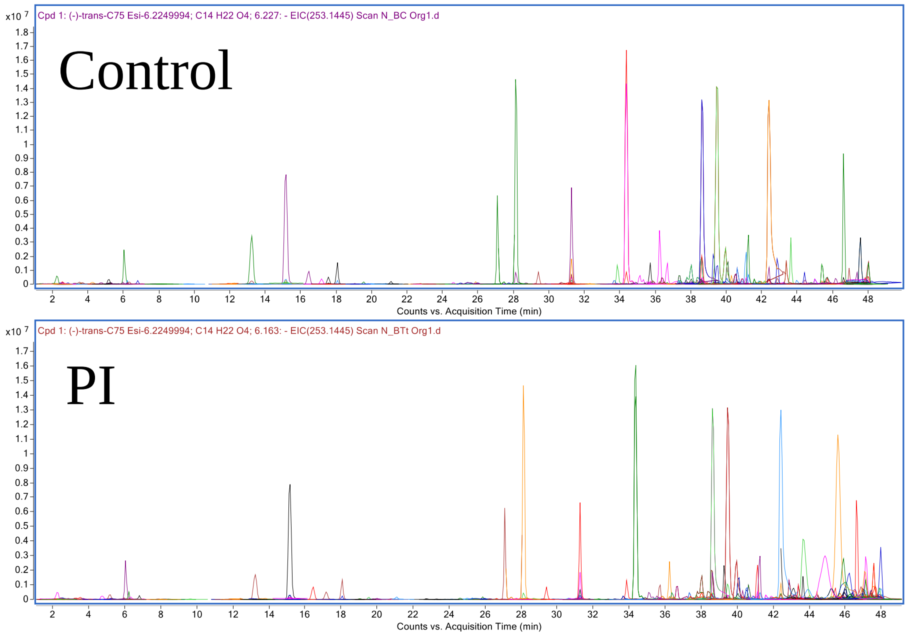

Metabolites recovered in the organic layer of the THP-1 cell extraction process under negative ion detection mode. Butanol: cells treated with n-butanol, the vehicle; PI: cells treated with PI

Neg_Org

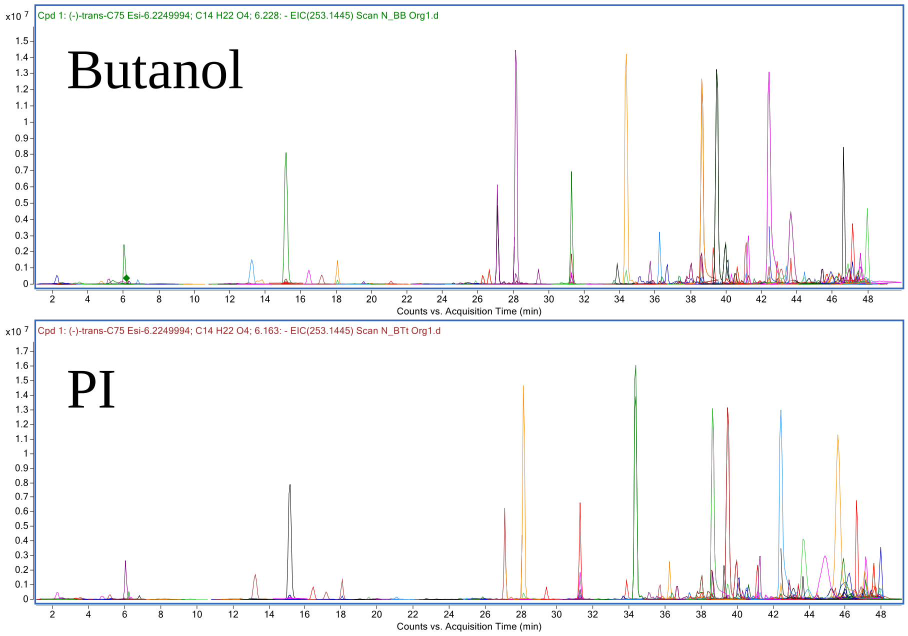

Metabolites recovered in the aqueous layer of the THP-1 cell extraction process under positive ion detection mode.  Control: untreated cell; PI: cells treated with PI

Pos_Aq

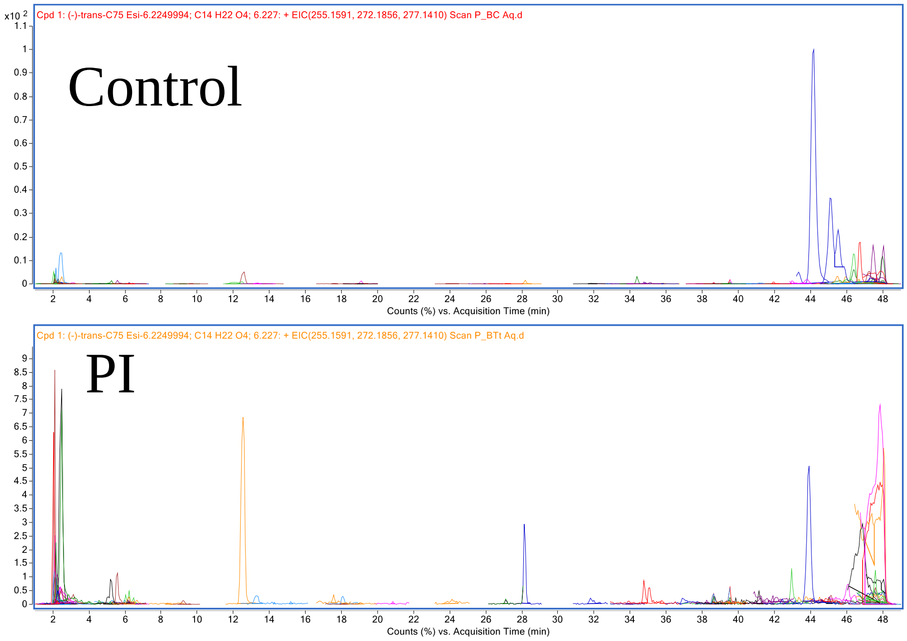

Metabolites recovered in the aqueous layer of the THP-1 cell extraction process under positive ion detection mode. Butanol: Cells treated with n-butanol, the vehicle; PI: cells treated with PI

Pos_Aq

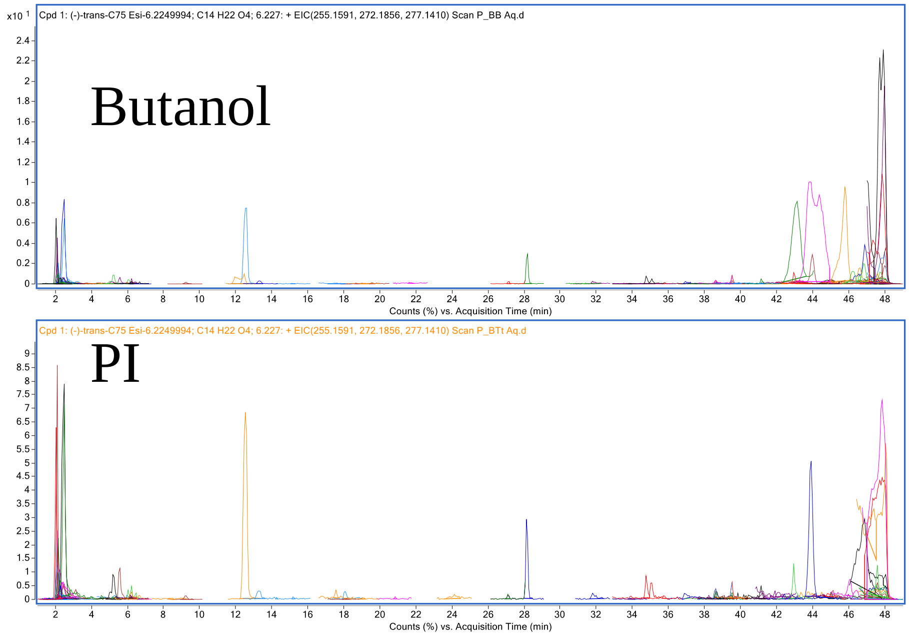

Metabolites recovered in the organic layer of the THP-1 cell extraction process under positive ion detection mode. Control: untreated cell; PI: cells treated with PI

Pos_Org

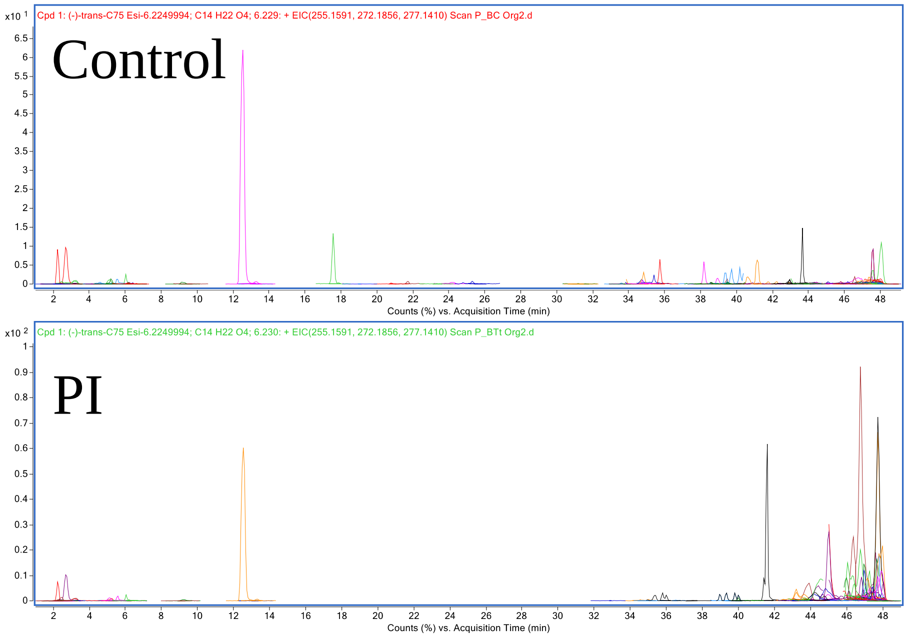

Metabolites recovered in the organic layer of the THP-1 cell extraction process under positive ion detection mode. Butanol: Cells treated with n-butanol, the vehicle; PI: cells treated with PI

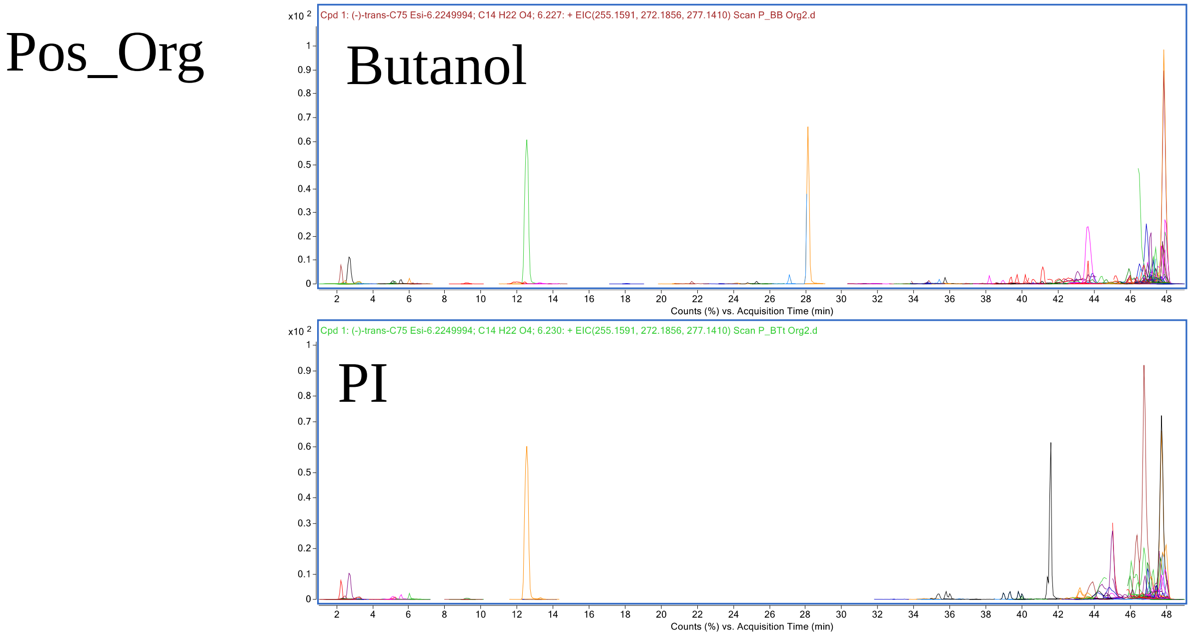
